# Solution structure of sperm lysin yields novel insights into molecular dynamics of rapid protein evolution

**DOI:** 10.1101/160580

**Authors:** Damien B. Wilburn, Lisa M. Tuttle, Rachel E. Klevit, Willie J. Swanson

**Affiliations:** Department of Genome Sciences, University of Washington, Seattle, WA, USA 98195; Department of Biochemistry, University of Washington, Seattle, WA, USA 98195

## Abstract

Protein evolution is driven by the sum of different physiochemical and genetic processes that usually results in strong purifying selection to maintain biochemical functions. However, proteins that are part of systems under arms race dynamics often evolve at unparalleled rates that can produce atypical biochemical properties. In the marine mollusk abalone, lysin and VERL are a pair of rapidly coevolving proteins that are essential for species-specific interactions between sperm and egg. Despite extensive biochemical characterization of lysin, including crystal structures of multiple orthologs, it was unclear how sites under positive selection may facilitate recognition of VERL. Using a combination of targeted mutagenesis and multidimensional NMR, we present a high-definition solution structure of sperm lysin from red abalone (*Haliotis rufescens*). Unapparent from the crystallography data, multiple NMR-based analyses conducted in solution reveal clustering of the N-and C-termini to form a nexus of 13 positively selected sites that constitute a VERL binding interface. Evolutionary rate was found to be a significant predictor of backbone flexibility, which may be critical for lysin bioactivity and / or accelerated evolution. These flexible, rapidly evolving segments that constitute the VERL binding interface were also the most distorted regions of the crystal structure relative to what was observed in solution. While lysin has been the subject of extensive biochemical and evolutionary analyses for more than 30 years, this study highlights the enhanced insights gained from applying NMR approaches to rapidly evolving proteins.

**Significance:** The fertilization of eggs by sperm is a critical biological process for nearly all sexually reproducing organisms to propagate their genetic information. Despite the importance of fertilization, the molecules that mediate egg-sperm interactions have been characterized for only a few species, and the biochemical mechanisms underlying these interactions are even less well understood. In the marine mollusk abalone, sperm lysin interacts with egg VERL in a species-specific manner to facilitate fertilization. In this report, we characterized the solution structure and molecular evolution of sperm lysin from red abalone (*Haliotis rufescens*), and identified the VERL binding interface as well as important lysin dimer interactions that have emerged as part of an incessant sexual arms race.

## Introduction

The innovation of elaborate male ornaments and female preferences by sexual selection has been of interest to evolutionary biologists since Darwin (1). Compared to other coevolving traits, sexual ornaments and preferences often evolve via runaway selection at exacerbated rates (2, 3). A similar phenomenon has been observed in that genes coding for reproductive proteins which evolve at extraordinary rates, and are usually among the fastest evolving genes in any genome (4). This pattern has been observed across diverse taxa (microbes, plants, and animals) and different types of reproductive proteins (sex pheromones, seminal fluid proteins, and gamete recognition proteins) (5). The rapid molecular evolution of reproductive genes is thought to be driven by the continual coevolution between interacting male and female protein pairs to optimize interactions. However, there presently exist few systems where both the interacting male and female proteins have been identified, creating a barrier to understanding the structural and biochemical interactions of paired reproductive proteins within an evolutionary framework.

A textbook example of such a rapidly evolving reproductive protein is abalone sperm lysin (6): a highly positively charged 16 kDa protein that is essential for dissolving the vitelline envelope (VE) of abalone oocytes (7). The VE – termed the zona pellucida (ZP) in mammals – is a glycoprotein envelope that serves as a barrier to the sperm and the external environment (8). In many taxa, sperm secrete proteins that non-enzymatically dissociate the VE, creating a hole through which sperm can pass, access the plasma membrane, fuse, and complete fertilization (4). While the molecular mechanisms and essential sperm proteins are unknown in mammals, the process is well characterized in abalone (illustrated in Figure 1). The abalone VE is largely composed of a gigantic ∼1 MDa glycoprotein, termed the <u>v</u>itelline <u>e</u>nvelope <u>r</u>eceptor for lysin (VERL), which contains 23 tandem repeats of a ZP-N polymerization domain. These ZP-N repeats are thought to polymerize through hydrogen bonds that form intermolecular β-sheets (9), and lysin dissolves the VE by competing for these hydrogen bonds (10). Molecular evolutionary analysis revealed that ∼25 of 134 residues are under positive selection (11), which may play a role in maintaining species boundaries. Off the North American Pacific coast, seven sympatric species of abalone are found with overlapping breeding seasons; while hybrids are viable in the lab, they are rare in nature (4). Additionally, lysin mediated VE dissolution is species – specific (10) and helps maintain species boundaries. Lysin is secreted as a dimer, suggesting an equivalent stoichiometry to bind dimerized VERL ZP-N repeats; however, prior biochemical experiments suggested the lysin monomer is the bioactive unit (12). The majority of the VERL repeats are >98% identical at the nucleotide level within a species, homogenized by concerted evolution (13). The two most N-terminal ZP-N repeats are not subjected to the same homogenization and are rapidly coevolving with lysin, suggestive of arms race dynamics (5, 14, 15). The specific molecular interactions between lysin and VERL that facilitate VE dissolution remain unknown.

**Figure 1.**
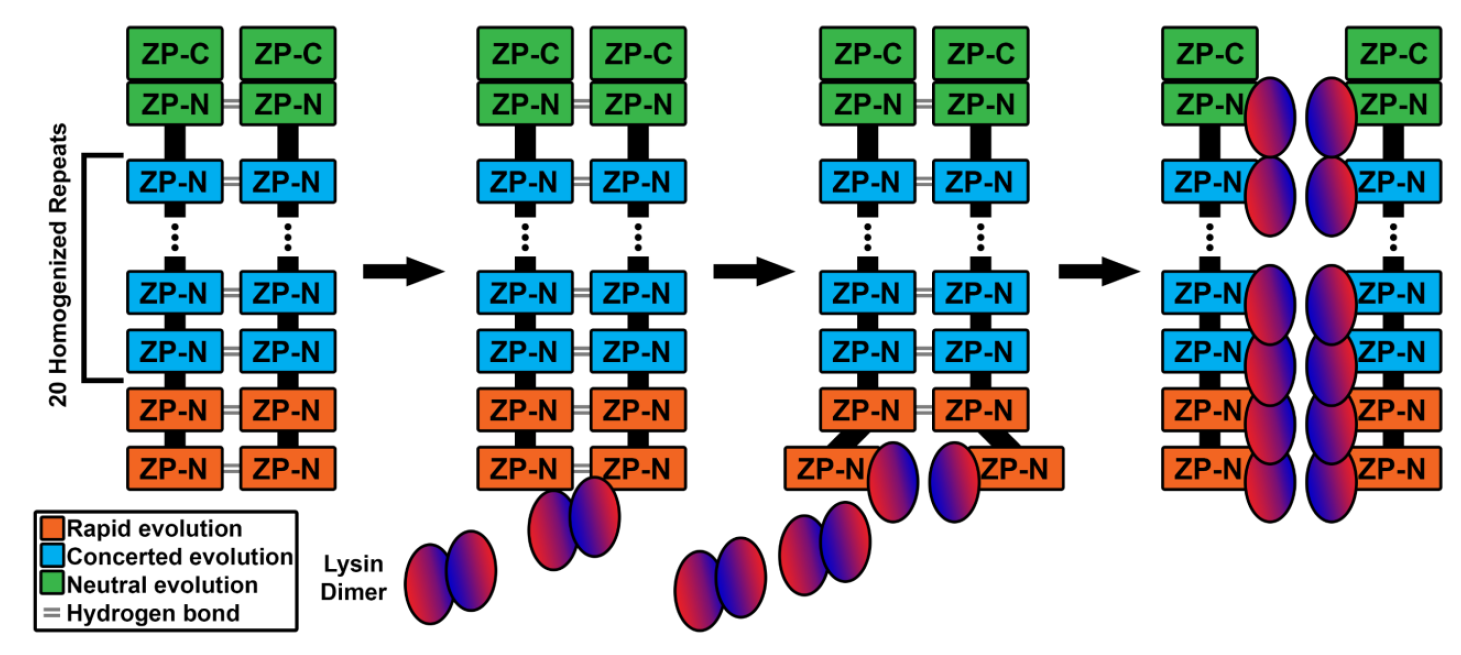
Proposed model of lysin-mediated VE dissolution. VERL is a gigantic molecule (∼1 MDa) that includes 23 repeats of a ZP-N polymerization module. The VE supramolecular structure is supported by intermolecular hydrogen bonds between VERL ZP-N repeats. The two most N-terminal repeats are rapidly coevolving with lysin while the remaining repeats are homogenized through concerted evolution (15). Upon sperm contact with the VE, lysin will disrupt the hydrogen bonds of the first repeat, forming a pair of lysin-VERL repeat heterodimers, and expose the next pair of repeats. Lysin can then disrupt the second pair of hydrogen-bonded VERL repeats, and this process continues until the VE is fully disrupted, allowing sperm to approach the egg plasma membrane.

Consistent with protein structure generally being more highly conserved than sequence (16), multiple structures determined by X-ray crystallography support that the structure of lysin is conserved, despite its extraordinarily rapid evolution. Nearly identical crystal structures were observed for lysin dimers of two divergent abalone species (*Haliotis rufescens*, red abalone, and *H. fulgens*, green abalone; backbone rmsd = 1.7 Å) (17, 18). A structure of monomeric red abalone lysin is highly similar to each dimeric subunit (average backbone rmsd = 1.0 Å) (18). However, the N-terminus is extended and highly unstructured (with no diffraction data for the first 3-4 residues) in all three structures, posing challenges for understanding lysin biology. The N-terminus harbors the largest concentration of positively selected sites (11), and mutagenesis experiments revealed that the N-terminus is critical for conferring species-specific recognition of the VE (19). Contacts formed between neighboring molecules in a protein crystal may disrupt or distort structure relative to the functional solution structure, thereby capturing structures that may not necessarily fully reflect biomolecules in their natural aqueous environments (20). To more fully assess the structural and dynamic features of rapidly evolving proteins, we used targeted mutagenesis and multidimensional NMR to determine the solution structure and subunit dynamics of sperm lysin.

## Results

### The lysin dimer is the bioactive form

The lysin dimer has modest affinity (KD ∼ 1 μM) and its subunits exchange rapidly. Based on dimer dissociation upon exposure to VEs, it was hypothesized that the monomer is the bioactive form (12). At concentrations required for NMR study (≥ 100 μM), lysin will exist almost exclusively as a dimer. As many of the 3D NMR experiments critical for structure determination are not sufficiently sensitive for proteins with molecular weights above ∼25 kDa, we developed a monomeric lysin mutant to use for in NMR-based analyses. Examination of the lysin dimer crystal structure revealed a set of three aromatic residues symmetrically mirrored across the dimer interface, with the phenylalanine 104 (F104) sidechain from each subunit in the center of the complex (Figure S1A). F104 is absolutely conserved in the seven California abalone species. Removal of this central aromatic ring by mutation of F104 to an alanine (F104A) produced a monomeric lysin without significantly altering the tertiary structure. Comparison of NMR spectra for WT and F104A lysin show only modest perturbations and these are near where the mutation was introduced (Figure S1B-C). The mean total rotational correlation time (τ_c_; which is proportional to molecular weight) for WT lysin (26.3 ns) was nearly twice that of F104A (13.7 ns, ratio = 1.92; Figure S1D), consistent with the mutation disrupting the dimer without causing a change in tertiary structure. Despite only perturbing the quaternary structure of the protein, F104A lysin was substantially less efficient at dissolving VEs compared to WT lysin (Figure 2A). Affinity pulldown of both WT and F104A lysin to recombinant VERL repeat 1 produced similar levels of binding relative to controls (Figure 2B), suggesting little or no difference in VERL affinity between dimeric and monomeric lysin. Therefore, while the lysin monomer can bind and recognize VERL repeats, the lysin dimer is required for efficient VE dissolution.

**Figure 2.**
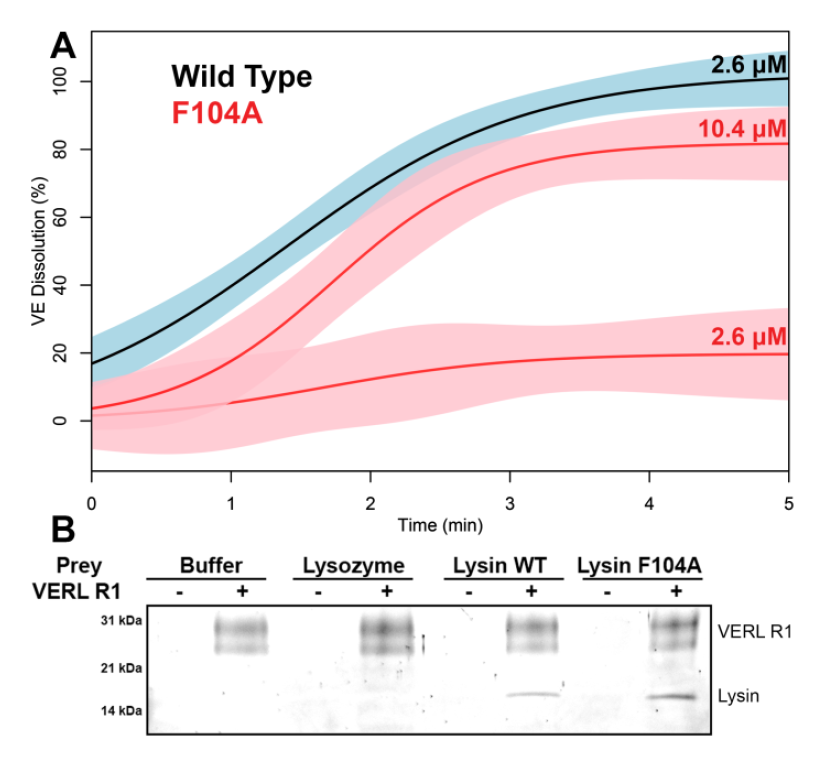
Monomeric lysin binds VERL, but is less efficient at VE dissolution. (A) *In vitro* VE dissolution experiments reveal that F104A lysin is ≥ 4-fold less efficient at dissolving VEs compared to WT, with lines of data fit to a logistic curve and shading of the 95% confidence interval. (B) Affinity pulldown experiments using recombinant VERL repeat 1 (R1) as bait show no major changes in binding affinity between WT and F104A lysin, suggesting that the effect of F104A on VE dissolution is a consequence of oligomeric state and not altered affinity for VERL.

### The VERL binding interface is comprised of several clustered positively selected sites

To understand how the N-terminus may be involved in interactions with individual VERL repeats – despite being “extended” in the available crystal structures – we determined the solution structure of F104A lysin using a suite of multidimensional heteronuclear NMR and restrained molecular dynamics simulation (Figure 3). An ensemble of the 20 lowest energy models was tightly constrained (backbone rmsd = 1.6 Å; Table S1), with regions of backbone flexibility at the N-terminus (residues 1-12) and in the turns that link α-helices (residues 39-43, 77-81, and 95-97). However, comparison of the solution and crystal structures revealed several discrepancies (Figure 4A-D). Most notably, the extended, disordered N-terminus in the crystal structure forms a nexus with the C-terminus and residues 60-65, driven largely by hydrophobic packing of the W3 sidechain with additional aromatic residues in this region (W62 and Y133; Figure S2). To validate the position of the N-terminus, we conducted experiments measuring the paramagnetic relaxation enhancement with a N-terminal Cu^2+^ binding tag (ATCUN; Figures 4C, S3A) and chemical shift perturbation analysis of a W3L mutant (Figures 4D, S3B). Neither the addition of the ATCUN tag nor the W3L mutation significantly impacted lysin VE dissolution activity (Figure S3C-D), suggesting that these mutations did not dramatically alter the tertiary structure and accurately reflect the coordination of the N-terminus with residues 60-65 and the C-terminus in solution. The second major discrepancy is with the α-helix that spans residues 44-76. In the crystal structure, there is a sharp turn at residues 61-63 not present in the NMR structure (Figure 4A), which likely reflects alternative hydrophobic packing due to the coordination of the N-terminus in this region.

**Figure 3.**
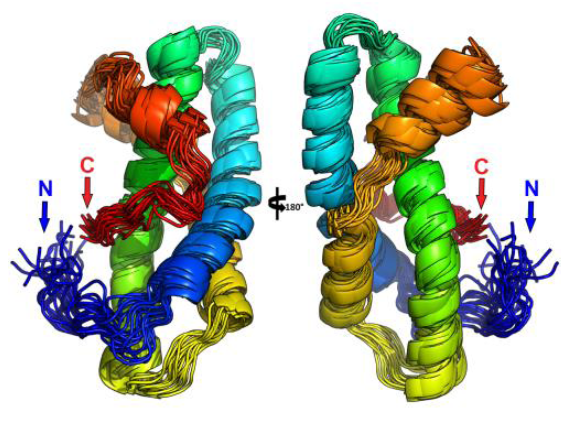
Ensemble of lysin solution structure. The ensemble of the 20 lowest energy structures of 100 simulated annealing simulations, colored in rainbow from N-terminus (blue) to C-terminus (red). The mean backbone rmsd for the constrained segment (residues 12-130) is 1.1 Å.

**Figure 4.**
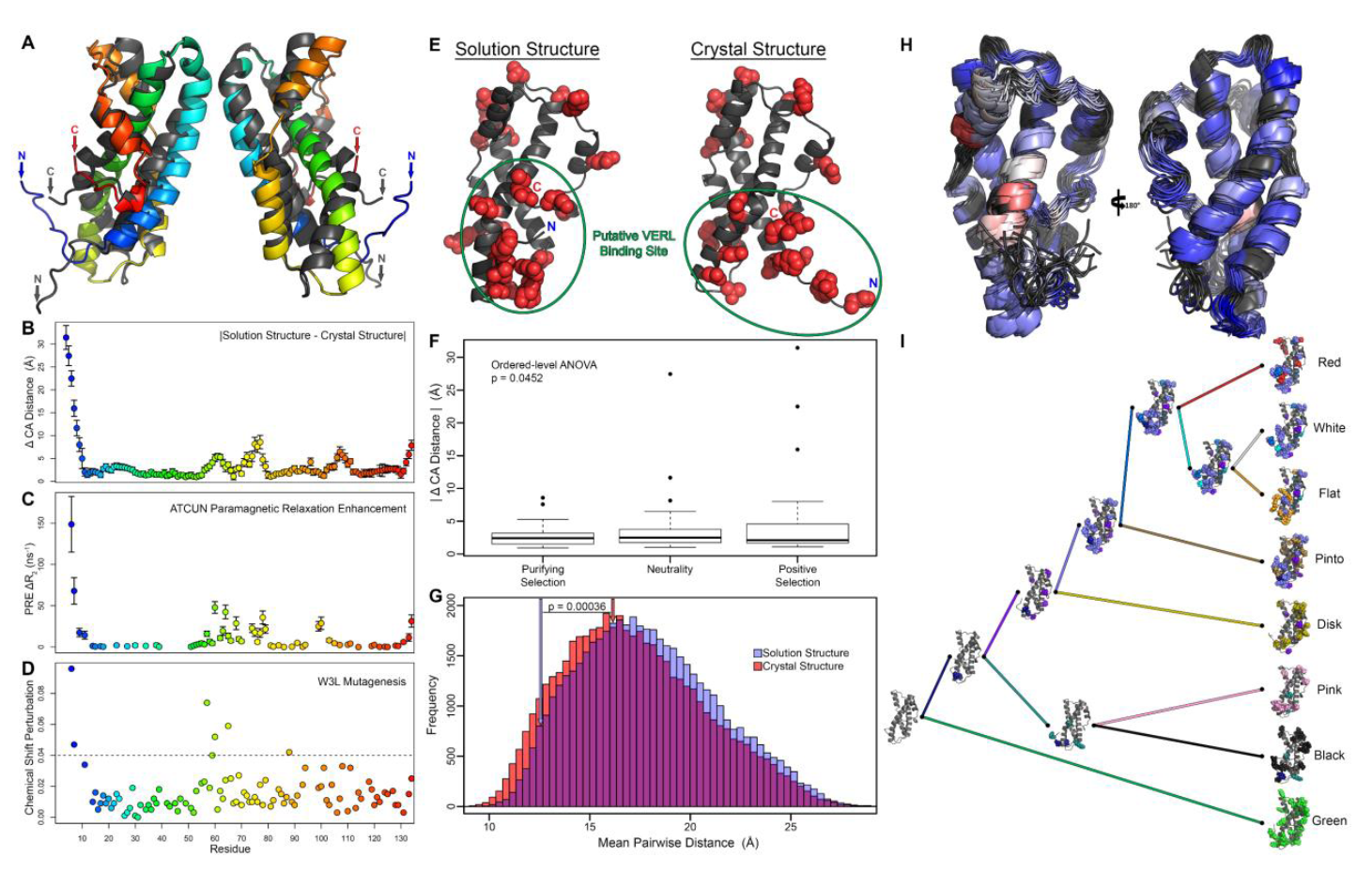
The VERL binding interface is composed of clustered positively selected sites unobserved in the crystal structure. (A) Structural alignment of the red abalone lysin crystal structure (PDB 2lis; grey) and the regularized average of the solution structure ensemble (rainbow). (B) Absolute value of the difference in CA positioning for each residue, plotted as the mean ± SD relative to each structure in the ensemble. The largest discrepancy is in the localization of the N-terminus. ATCUN-based paramagnetic relaxation enhancement (C) and chemical shift perturbation from a W3L mutation (D) provides independent support that the N-terminus clusters near residues 60-65 and the C-terminus. (E) Comparison of the solution and crystal structures of lysin, with side chains of positively selected sites highlighted as red spheres and the putative VERL binding site circled in green. (F) Comparison of residue positions between the solution and crystal structures divided by mode of evolution (purifying selection, neutrality, or positive selection based on the M2A model), demonstrating a position correlation between structural discrepancy and dN / dS. (G) Thirteen positively selected sites found at the nexus between the N-terminus, C-terminus, and residues 64-79 (circled in green in panel A) cluster better than 95% of similarly spaced residues, and likely constitute the VERL recognition site., The clustering of these residues (with arrows highlighting the value for the 13 important positively selected sites) was lost in the crystal structure, and the difference between the solution and crystal structures is significant (p = 0.00036). (H) Titration of VERL repeat 1 against NMR-reactive F104A lysin confirms that the region of clustered positively selected sites is the VERL interface. The ensemble structure is colored based on the backbone amide relative signal intensity for each residue which reflects changes in chemical shift and / or relaxation upon VERL binding, colored from blue to white to red corresponding to increasing perturbation. (I) Homology modeling of extant and ancestral states of lysin sequences, with mutations acquired on each branch individually colored. There is continual replacement of the amino acids at the VERL binding interface, consistent with incessant evolution of this region.

Positively selected sites of binding proteins are often clustered near the ligand binding domain to maintain high affinity interactions (21-23). A similar clustering on lysin would be expected to define the VERL recognition face. Comparison of the solution and crystal structures revealed dramatic differences in the packing of positively selected sites (Figure 4E), and there was a significant positive correlation between evolutionary rate (dN / dS) and the magnitude by which the crystal structure deviated from the solution structure ensemble (Figure 4F). While there are positively selected sites widely distributed across the surface of lysin, the convergence of the N- and C-termini with residues 64-79 results in the clustering of 13 positively selected residues. When compared to a null distribution of similarly spaced residues, these 13 residues clustered better than >95% of cases (mean pairwise CA distance = 12.6 Å). In the crystal structure, these same 13 sites were significantly further apart (16.1 Å, p = 0.00036 Figure 4G). Given the concentration of positively selected sites, and that the N- and C-termini are essential for species-specific VE dissolution (19), we hypothesized that this region is the principal interface for lysin-VERL interactions. To biochemically test this hypothesis, NMR perturbation analysis was performed and unlabeled red VERL repeat 1 was added incrementally to ^15^N-labeled F104A red lysin until the two proteins were at equimolar concentrations, with points of contact measured by changes in peak intensity (Figure S4). Three residues (T60, A63, and Y100) were particularly sensitive, and were undetectable after addition of 30% VERL repeat 1, suggesting significant changes in backbone chemical shift and/or exchange rates upon binding to VERL (Figure 4H). All three residues are proximal to the cluster of rapidly evolving sites, validating this region as the VERL binding interface. Using homology modeling, structures of extant and ancestral states of lysin were constructed to understand the process of mutation acquisition across Pacific abalone. In a phylogenetic context, the VERL binding interface shows repeated mutation acquisition that is consistent with lysin likely having an important role in maintaining species-specific interactions and providing a barrier to hybridization (Figure 4I).

### Molecular dynamics of lysin correlate with evolutionary rate

To our knowledge, lysin is the first rapidly evolving reproductive protein for which both crystal and solution structures have been solved. Given that the discrepancies between the two structures were correlated with rate of evolution (Figure 4F), we sought to further address how strong directional selection from a coevolving partner may influence various aspects of protein structure. First, a significant advantage of NMR over X-ray crystallography is that protein dynamics can be measured through NMR relaxation analysis. Backbone amide measurements of τc and heteronuclear NOE (two measurements of different nuclear relaxation properties) both reveal positive correlation between structural flexibility and dN/dS (Figure S5B). These more flexible residues are primarily located on the termini and in the loops between the α-helices (Figure S5A), but these were also the regions that were the least accurately reflected in the crystal structure (Figure 4B). The relationship between structural flexibility and positively selected sites inspired us to consider how evolutionary rate might influence intra-residue covariation, as multiple groups have found a negative correlation between covariation and structural distance, likely due to epistasis between neighboring sites to maintain protein folding (24, 25). If rapidly evolving sites are on flexible segments that are largely subjected to coevolution with VERL, then there is likely reduced epistasis from other lysin residues and we would expect no relationship between covariation and structural distance. We approximated intra-residue covariation by inferring ancestral states at each node in the lysin gene tree using maximum likelihood, and counted how frequently two sites change on the same branch or not (normalized against the mean number of times each site changes). Structural distance, evolutionary rate, and their interaction term were all significant predictors of intra-residue covariation (2-way ANOVA, p < 10^-4^ for all variables). While there is a significant negative correlation between covariation and structural distance for sites either jointly under purifying selection or neutrality, there is no relationship for sites jointly under positive selection (Figure S6). While there was only measurable covariation for a few pairs of sites jointly under purifying selection, it is noteworthy that these pairs had a higher mean covariation (higher y-intercept), stronger relationship (sharper slope), and were the only examples of 100% covariation. Hence, evolutionary rate – and likely the source of the evolutionary pressure – is a significant corollary of intra-residue covariation.

## Discussion

Fertilization – an essential biological process for all sexually reproducing taxa – has been the subject of a century of ongoing research, yet despite its obvious biological importance, characterizing the underlying molecular mechanisms has remained a challenge. A key confounder in this research is rapid protein evolution: while the major cellular events of fertilization are broadly conserved across eukaryotes, the molecules mediating the processes diverge rapidly (4). Abalone sperm lysin was purified 30 years ago (7), and is still one the few fertilization proteins for which its receptor has been identified (10, 26, 27). It remains a critical model both for characterizing the mechanisms of VE dissolution and for understanding how pervasive sexual selection from a coevolving receptor can shape protein structure. In this study, we have shed new insights into the evolution of structure, function, and dynamics of this classic reproductive protein.

We generated monomeric lysin via a single point substitution of a residue at the dimer interface (F104A). The mutation causes minimal perturbation to the tertiary structure (Figure S1), maintains affinity for VERL, but decreases VE dissolution activity (Figure 2), suggesting that oligomer state is important for lysin’s function (12). While monomeric F104A lysin is less efficient, it is still capable of dissolving VEs at sufficiently high concentrations (Figure 2). We hypothesize that dissolution involves a competition between the binding of lysin to a VERL repeat to form a lysin-VERL heterodimer and binding of another VERL repeat to a VERL repeat to form a VERL repeat homodimer (Figure S7). Although the monomeric F104A-lysin can bind, it is less effective at competing with another VERL repeat that is in high local concentration in the VE. In contrast, binding of a subunit of dimeric lysin to a VERL repeat effectively increases the local concentration of unbound lysin subunit (i.e., the second subunit in the dimer) to allow its simultaneous binding to the neighboring VERL repeat. This concentration dependence may become more important as the VE begins to dissolve – with the next pair of exposed VERL repeats deeper in the dissolution furrow, the simultaneous diffusion of two lysin monomers into this space becomes probabilistically infeasible. The lysin dimer may be a compensatory structural feature to overcome the expanded ZP-N array of VERL. Based on *in silico* docking simulations using lysin and VERL paralogs, our data suggest that lysin transitioned from a monomer to a dimer near the same evolutionary point that the VERL ZP-N array expanded (Figure 5). Duplication of N-terminal ZP-N repeats is common in many egg coat proteins (28), but the functional significance has remained unclear. Field experiments in sea urchin suggest that under sufficiently high population densities, polyspermy risk can drive selection of oocytes with weaker affinity to sperm, spurring an arms race and pulling the co-evolving egg-sperm proteins across the phenotypic landscape (29, 30). In an ancestral abalone population, similar polyspermy risks may have prompted duplication of the VERL ZP-N array, increasing the required lysin concentration to dissolve the VE and eventually necessitating the transition from monomer to dimer.

**Figure 5.**
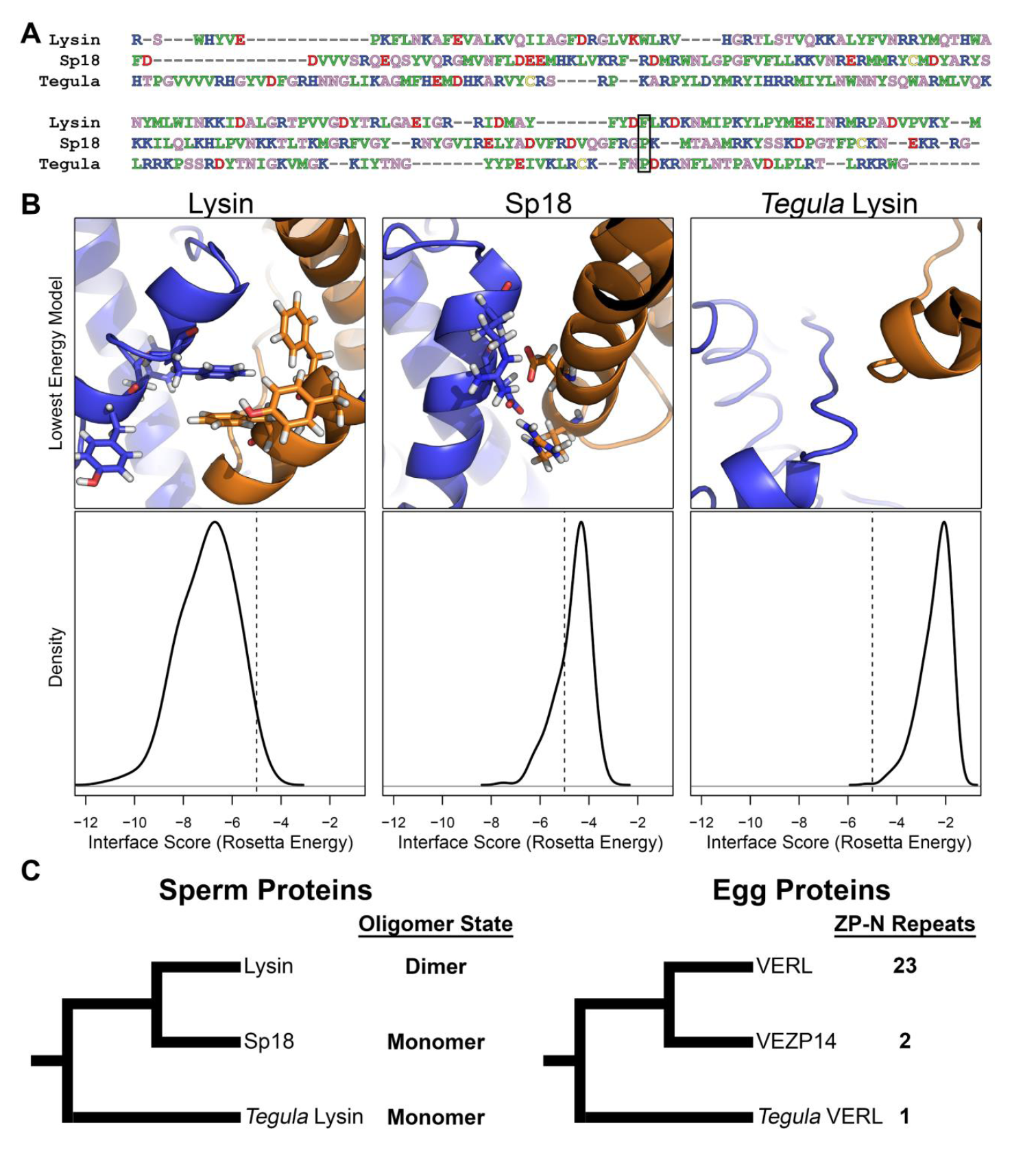
The lysin dimer is a compensatory mutation to overcome the expansion of the VERL ZP-N array. To characterize the evolutionary history of lysin and its oligomeric state, we compared its sequence and structure with that of abalone Sp18 and black tegula lysin. (A) Sequence alignment of the three proteins reveals that the critical F104 for dimer formation (highlighted with a black box) is a proline in both Sp18 and *Tegula* lysin. (B) Rosetta-based *in silico* docking simulations was performed to infer the oligomer state of each protein. From 10,000 simulations, the interface score density was compared for the 5% lowest energy structures, with values below-5 suggestive of strong docking (48). The median structure of these low-energy interfaces is shown. In lysin model, the essential aromatic triad centered around F104 is recapitulated at the interface; in Sp18, the homologous region is replaced with charged residues that were predicted to form two weak salt bridges and in *Tegula* lysin, there is no significant interface. (C) Our combined sequence and structural analysis suggest that the lysin dimer is the derived state in abalone, and comparison of ZP-N repeats within VERL homologs (abalone VEZP14 and *Tegula* VERL) suggest that the lysin dimer arose near the same evolutionary point as when the VERL ZP-N array expanded. It should be noted that the specific interacting partners of Sp18 and VEZP14 are currently unknown (49, 50).

The solution structure of lysin presented here also reveals another key aspect of its role in fertilization: the VERL binding interface, formed by the spatial clustering of positively selected sites that are perturbed upon binding by VERL (Figure 4E,H). The clustering of these sites reflects an interesting interplay between positive selection and structural flexibility. As accurate modeling of flexible regions poses a challenge to crystallographic studies (20), reliance on crystal structures for insights pertaining to region under high positive selection may limit the insights provided for rapidly evolving proteins. The correlation between evolutionary rate and structural flexibility drives a “chicken and egg” quandary – “which comes first, flexibility or rapid evolution?” From a genetics perspective, rapid evolution may facilitate structural flexibility: as arms race dynamics with a coevolving partner promotes acquisition of non-synonymous substitutions, epistasis between residue positions begins to decompose and the segment becomes more flexible. However, re-framing the question from a structural perspective likely presents the opposing direction: fewer physiochemical constraints in regions of the protein allows greater tolerance for non-synonymous substitutions, such that they may be more frequently co-opted as recognition faces because of relaxed purifying selection. Unfortunately, there are few studies that have explicitly examined the relationship between flexibility and rapid evolution. One of the other few reproductive proteins for which an NMR structure has been determined is Plethodontid Modulating Factor (PMF), a protein pheromone secreted by male plethodontid salamanders to regulate female courtship behavior (31). The structure of PMF is largely stabilized by four disulfide bonds, a hallmark of the three-finger protein superfamily of which PMF is a member; however, PMF has an alternate disulfide bonding pattern compared to other members that reduces its secondary structure and creates flexible loops with several rapidly evolving sites. These conserved disulfide bonds may be analogous to the hydrophobic interactions of W3-W62-Y134 and act as “anchor points” to allow neighboring loops to evolve with reduced purifying selection. While this suggests both reproductive systems likely favor the structural “flexibility first” hypothesis, it is easy to envision how natural purifying selection could favor highly stabilizing interactions (disulfide bridges and / or π-π overlap from aromatic residues) as a compensatory process to handle reduced epistasis resulting from strong positive sexual selection. The role of flexibility in adaptive coevolution of rapidly evolving proteins remains a subject that warrants greater investigation, by using NMR and / or other solution-based biophysical techniques in conjunction with evolutionary studies.

## Conclusions

Reproductive proteins are a classic example of rapid protein evolution, with differing reproductive strategies between sexes driving arms race dynamics and accelerated coevolution between interacting proteins. Abalone sperm lysin is arguably the most well studied reproductive protein, and through adaptive coevolution with its egg coat receptor VERL, lysin has experienced repeated bursts of positive selection to maintain high affinity interactions and possibly to maintain species boundaries. In this report, we provide new insights into the function of this classic evolutionary system by determining the solution structure of lysin. Functionally, we determined that lysin oligomerization plays a central role in egg coat dissolution, likely as an evolutionary consequence of sexual conflict and duplication of ZP-N repeats within VERL. Structurally, our solution structure revealed clustering of positively selected residues that form the essential VERL binding interface. Notably, the same clustering was not observed in the crystal structure, and observed correlations between structural flexibility and evolutionary rate warrant greater consideration when exploring the structures of rapidly evolving proteins. While lysin has been the subject of extensive biochemical characterization for more than 30 years, our findings shed new insights into the function and adaptive coevolution of this classic fertilization protein.

## Methods

See Supplemental Methods for complete details.

### Cloning and expression of lysin

Methods for lysin cloning, expression, and purification were adapted from Lyon and Vacquier (19). The lysin coding sequence was PCR amplified from *H. rufescens* testis cDNA and cloned into the pET11d vector (Promega, Madison, WI). Lysin mutants (F104A, W3L, F104A/W3L, and addition of ATCUN motif) were engineered into the plasmid using the Q5 site-directed mutagenesis kit (New England Biolabs). All lysin constructs were transformed into Rosetta2gami chemically competent *E. coli* (EMD-Millipore, Billerica, MA) and validated by Sanger sequencing (Eurofin Genomics, Louisville, KY). Lysin was expressed as insoluble inclusion bodies, solubilized with 5M guanidine-HCl in buffered sea water, refolded by rapid dilution, and purified by cation exchange chromatography. Isotopic labeling for NMR experiments was performed using methods adapted from Marley et al. (32).

### Lysin functional analysis

VE dissolution assays were performed using methods adapted from Lewis et al. (7). Abalone VEs were isolated from gravid female abalone by dissection of ovaries, and oocytes lysed by detergent treatment (1% Triton X-100 in buffered sea water). VE dissolution was performed by aliquoting 100 μL of VEs, adding 20 μL of lysin (containing 5 or 20 μg), measuring the absorbance at 640 nm (turbidity) every 30 seconds, and calculating the percent dissolution. All VE dissolution experiments were repeated 6 times, and data fit via nonlinear regression to a logistic curve using the R function nlsLM from the minpack.lm package. For VERL pulldown analysis, *H. rufescens* VERL repeat 1 was cloned and expressed in *Pichia pastoris* (och1^-^,aox1^-^). An N-terminal Streptavidin Binding Peptide (SBP-tag) was included for joint purification and affinity pulldown. Lysin pulldown experiments were performed loading streptavidin magnetic beads (New England Biolabs) with ∼5 μg recombinant VERL repeat 1, equilibrating the beads with buffered sea water, incubating the beads with buffer or 20 μg prey (lysozyme or a lysin variant) in 60 μL buffered sea water with 1 mg / mL BSA for 30 minutes, rinsing the beads three times with buffered sea water, transferring the beads to a clean microfuge tube, and eluting the VERL-lysin complex with 20 μL 0.02% biotin/0.5M NaCl / 20mM Tris, pH 8. Elutions were analyzed using Tris-Tricine SDS-PAGE (33), and gels stained using SYPRO Ruby (Invitrogen).

### NMR analysis of recombinant lysin

Purified, isotopically-labeled lysin (^15^N or ^15^N / ^13^C) was concentrated to ∼150-450 μM in 200mM NaCl / 10mM NaPO_4_, pH 7 / 7% D_2_O, and all NMR experiments were performed on either a Bruker Avance 500 MHz spectrometer or on a Bruker Avance 600 MHz spectrometer fitted with a TCI CryoProbe (Bruker, Billerica, MA). NMR assignments were obtained using a combination of standard 2D / 3D experiments, with spectra analyzed using NMRFAM-SPARKY (34), and initial structure calculations performed with CYANA 2.1 (35, 36) using automatic NOE assignments and dihedral angle restraints (from TALOS-N (37) or measured ^3^JHN-HA couplings) CYANA structures were refined using Xplor-NIH 2.43 (38, 39) with residual dipolar couplings (40) and explicit solvation (eefx) (41), with simulated annealing from 3500 to 25 K followed by Cartesian minimization. Spin-lattice (longitudinal) relaxation constants (R_1_), spin-spin (transverse) relaxation rate constants (R_2_), and ^15^N[^1^H] steady-state heteronuclear NOEs of the backbone ^15^N nuclei were measured at 500 Mhz and 298 K. Position of the lysin N-terminus was validated by ATCUN PRE measurements (42, 43). To characterize the VERL binding interface, ^15^N-HSQC spectra of F104A lysin were acquired with different concentrations of VERL repeat 1, with sensitivity to VERL titration reported as normalized peak intensities. All NMR data were deposited in the BMRB (30246) and the structural ensemble deposited in the RCSB PDB (5UTG).

### Structural Analysis

Lysin sequences from 29 abalone species were collected from Genbank, aligned using FSA (44), gene tree was constructed with amino acid sequences using RAxML and the JTT substitution model (45), and evolutionary rates calculated using PAML 4.8 (46). Sites under positive selection are defined as having a Naïve Empirical Bayes (NEB) probability of greater than 0.9 under the M8 model. For comparison of sites under purifying, neutral, or positive selection, residues are grouped based on the class with the highest NEB posterior probability under the M2A model. Using ancestral states constructed by PAML on each node of the lysin gene tree, intra-residue covariation was calculated as the ratio of the number of times two residues change on the same branch divided by the mean number of times both residues change total. Weighted linear regression was performed to compare the relationship between covariation, mean CA distance, and evolutionary rate, with weights calculated as the product of the posterior probabilities for each ancestral state being applied. Docking simulations were performed using Rosetta 3.5 (47, 48), with templates including the lysin F104A solution structure (*in silico* mutagenized back to WT), the Sp18 crystal structure, and a homology model of *Tegula funebralis* lysin constructed using Rosetta. Each template structure was energy minimized in Rosetta using the relax function, structures duplicated and aligned to each subunit in the red lysin dimer crystal structure (PDB 2lyn), and 10,000 docking simulations were performed. The interface score of the top 5% lowest energy structures were analyzed, with values below-5 considered a good docking score (48).

## Acknowledgements

We thank Dr. Jan Aaagaard and Emily Killingbeck for comments on the manuscript. This work was supported by National Institutes of Health grants R01-HD076862 to W.J.S., F32-GM116298 to D.B.W.

## References

Darwin CR (1871) The descent of man, and selection in relation to sex (John Murray, UK).

Lande R (1981) Models of speciation by sexual selection on polygenic traits. Proceedings of the National Academy of Sciences 78(6):3721&3725.

Mead LS & Arnold SJ (2004) Quantitative genetic models of sexual selection. Trends in Ecology & Evolution 19(5):264&271.

Swanson WJ & Vacquier VD (2002) The rapid evolution of reproductive proteins. Nature Review Genetics 3(2):137&144.

Wilburn DB & Swanson WJ (2016) From molecules to mating: Rapid evolution and biochemical studies of reproductive proteins. Journal of Proteomics 135:12&25.

Barton NH, Briggs DEG, Eisen JA, Goldstein DB, & Patel NH (2007) *Evolution* (Cold Spring Harbor Laboratory Press).

Lewis CA, Talbot CF, & Vacquier VD (1982) A protein from abalone sperm dissolves the egg vitelline layer by a nonenzymatic mechanism. Developmental Biology 92(1):227&239.

Bleil JD & Wassarman PM (1980) Structure and function of the zona pellucida: identirication and characterization of the proteins of the mouse oocyte’s zona pellucida. Developmental Biology 76(1):185&202.

Bokhove M, et al. (2016) A structured interdomain linker directs self-polymerization of human uromodulin. Proceedings of the National Academy of Sciences 113(6):1552&1557.

Swanson WJ & Vacquier VD (1997) The abalone egg vitelline envelope receptor for sperm lysin is a giant multivalent molecule. Proceedings of the National Academy of Sciences 94(13):6724&6729.

Yang Z, Swanson WJ, & Vacquier VD (2000) Maximum-likelihood analysis of molecular adaptation in abalone sperm lysin reveals variable selective pressures among lineages and sites. Molecular Biology and Evolution 17(10):1446&1455.

Shaw A, Fortes PA, Stout CD, & Vacquier VD (1995) Crystal structure and subunit dynamics of the abalone sperm lysin dimer: egg envelopes dissociate dimers, the monomer is the active species. The Journal of Cell Biology 130(5):1117&1125.

Swanson WJ & Vacquier VD (1998) Concerted evolution in an egg receptor for a rapidly evolving abalone sperm protein. Science 281(5377):710&712.

Galindo BE, Moy GW, Swanson WJ, & Vacquier VD (2002) Full-length sequence of VERL, the egg vitelline envelope receptor for abalone sperm lysin. Gene 288:111&117.

Clark NL, et al. (2009) Coevolution of interacting fertilization proteins. PLOS Genetics

5(7):e1000570.

Moult J, Fidelis K, Kryshtafovych A, Schwede T, & Tramontano A (2016) Critical assessment of methods of protein structure prediction: Progress and new directions in round XI. Proteins: Structure, Function, and Bioinformatics 84:4&14.

Kresge N, Vacquier VD, & Stout CD (2000) The high resolution crystal structure of green abalone sperm lysin: implications for species-specific binding of the egg receptor. Journal of Molecular Biology 296:1225&1234.

Kresge N, Vacquier VD, & Stout CD (2000) 1.35 and 2.07 A resolution structures of the red abalone sperm lysin monomer and dimer reveal features involved in receptor binding. Acta Crystallographica Section D 56(1):34&41.

Lyon JD & Vacquier VD (1999) Interspecies chimeric sperm lysins identify regions mediating species-specific recognition of the abalone egg vitelline envelope. Developmental Biology 214:151&159.

Rupp B (2009) Biomolecular Crystallography: Principles, Practices, and Applications to Structural Biology (Garland Science) 1st edition Ed.

Nei M, Gu X, & Sitnikova T (1997) Evolution by the birth-and-death process in multigene families of the vertebrate immune?system. Proceedings of the National Academy of Sciences 94(15):7799&7806.

da Fonseca RR, Kosiol C, Vinař T, Siepel A, & Nielsen R (2010) Positive selection on apoptosis related genes. FEBS Letters 584(3):469&476.

Clark NL & Swanson WJ (2005) Pervasive adaptive evolution in primate seminal proteins. PLOS GENETICS 1(3):e35.

Morcos F, et al. (2011) Direct-coupling analysis of residue coevolution captures native contacts across many protein families. Proceedings of the National Academy of Sciences 108(49):E1293&E1301.

Hopf TA, et al. (2014) Sequence co-evolution gives 3D contacts and structures of protein complexes. eLife 3:e03430.

Hart MW (2013) Structure and evolution of the sea star egg receptor for sperm bindin. Molecular Ecology 22(8):2143&2156.

Bianchi E, Doe B, Goulding D, & Wright GJ (2014) Juno is the egg Izumo receptor and is essential for mammalian fertilization. Nature 508(7497):483&487.

Callebaut I, Mornon JP, & Monget P (2007) Isolated ZP-N domains constitute the N-terminal extensions of Zona Pellucida proteins. Bioinformatics 23(15):1871&1874.

Levitan DR & Stapper AP (2010) Simultaneous positive and negative frequency-dependent selection on sperm bindin, a gamete recognition protein in the sea urchin *Strongylocentrotus purpuratus*. Evolution 64(3):785&797.

Stapper AP, Beerli P, & Levitan DR (2015) Assortative mating drives linkage-disequilibrium between sperm and egg recognition protein loci in the sea urchin *Strongylocentrotus purpuratus*. Molecular Biology and Evolution 32(4):859&870.

Wilburn DB, et al. (2015) Pheromone isoform composition differentially affects female behaviour in the red-legged salamander, *Plethodon shermani*. Animal Behaviour 100:1&7.

Marley J, Lu M, & Bracken C (2001) A method for efficient isotopic labeling of recombinant proteins. Journal of Biomolecular NMR 20:71&75.

Schägger H & von Jagow G (1987) Tricine-sodium dodecyl sulfate-polyacrylamide gel electrophoresis for the separation of proteins in the range from 1 to 100 kDa. Analytical Biochemistry 166:368&379.

Lee W, Tonelli M, & Markley JL (2015) NMRFAM-SPARKY: enhanced software for biomolecular NMR spectroscopy. Bioinformatics 31(8):Epub 2014 Dec 2012.

Güntert P, Mumenthaler C, & Wüthrich K (1997) Torsion angle dynamics for NMR structure calculation with the new program Dyana. Journal of Molecular Biology 273(1):283&298.

Herrmann T, Güntert P, & Wüthrich K (2002) Protein NMR structure determination with automated NOE assignment using the new software CANDID and the torsion angle dynamics algorithm DYANA. Journal of Molecular Biology 319(1):209&227.

Shen Y & Bax A (2013) Protein backbone and sidechain torsion angles predicted from NMR chemical shifts using artificial neural networks. Journal of Biomolecular NMR 56:227&241.

Schwieters CD, Kuszewski JJ, Tjandra N, & Clore GM (2003) The Xplor-NIH NMR Molecular Structure Determination Package. Journal of Magnetic Resonance 160:66&74.

Schwieters CD, Kuszewski JJ, & Clore GM (2006) Using Xplor-NIH for NMR molecular structure determination. Progress in Nuclear Magnetic Resonance Spectroscopy 48:47&62.

Higman VA, Boyd J, Smith LJ, & Redfield C (2011) Residual dipolar couplings: are multiple independent alignments always possible? Journal of Biomolecular NMR 49(1):53&60.

Tian Y, Schwieters CD, Opella SJ, & Marassi FM (2014) A practical implicit solvent potential for NMR structure calculation. Journal of Magnetic Resonance 243:54&64.

Donaldson LW, et al. (2001) Structural characterization of proteins with an attached ATCUN motif by paramagnetic relaxation enhancement NMR spectroscopy. Journal of American Chemical Society 123:9843&9847.

Lee GM, et al. (2008) The affinity of Ets-1 for DNA is modulated by phosphorylation through transient interactions of an unstructured region. Journal of Molecular Biology 382(4):1014&1030.

Bradley RK, et al. (2009) Fast Statistical Alignment. PLOS Computational Biology 5:e1000392.

Stamatakis A (2014) RAxML Version 8:A tool for Phylogenetic Analysis and Post-Analysis of Large Phylogenies. Bioinformatics 30(9):1312&1313.

Yang Z (2007) PAML 4: Phylogenetic Analysis by Maximum Likelihood. Molecular Biology and Evolution 24(8):1586&1591.

Gray JJ, et al. (2003) Protein-protein docking with simultaneous optimization of rigid-body displacement and side-chain conformations. Journal of Molecular Biology 331(1):281&299.

Chaudhury S, et al. (2011) Benchmarking and analysis of protein docking performance in RosettaDock 3.2. PLOS ONE 6(8):e22477.

Aagaard JE, Vacquier VD, MacCoss MJ, & Swanson WJ (2010) ZP domain proteins in the abalone egg coat include a paralog of VERL under positive selection that binds lysin and 18-kDa sperm proteins. Molecular biology and evolution 27(1):193&203.

Aagaard JE, Springer SA, Soelberg SD, & Swanson WJ (2013) Duplicate abalone egg coat proteins bind sperm lysin similarly, but evolve oppositely, consistent with molecular mimicry at fertilization. PLoS Genet 9(2):e1003287.

Liang S, Zou C, Lin Y, Zhang X, & Ye Y (2013) Identification and characterization of PGCW14: a novel strong constitutive promoter of *Pichia pastoris*. Biotechnology Letters 35:1865&1871.

Liachko I & Dunham MJ (2014) An autonomously replicating sequence for use in a wide range of budding yeasts. FEMS yeast research 14(2):364&367.

Wu S & Letchworth GJ (2004) High efficiency transformation by electroporation of *Pichia pastoris* pretreated with lithium acetate and dithiothreitol. BioTechniques 36:152&154.

